# Macrophage Scavenger Receptor 1 mediates lipid-induced inflammation in non-alcoholic fatty liver disease

**DOI:** 10.1101/2020.02.01.930115

**Authors:** Olivier Govaere, Sine Kragh Petersen, Nuria Martinez-Lopez, Jasper Wouters, Matthias Van Haele, Rosellina M. Mancina, Oveis Jamialahmadi, Orsolya Bilkei-Gorzo, Pierre Bel Lassen, Rebecca Darlay, Julien Peltier, Jeremy M. Palmer, Ramy Younes, Dina Tiniakos, Guruprasad P. Aithal, Michael Allison, Michele Vacca, Melker Göransson, Michael J Drinnan, Hannele Yki-Järvinen, Jean-Francois Dufour, Mattias Ekstedt, Sven Francque, Salvatore Petta, Elisabetta Bugianesi, Jörn M Schattenberg, Christopher P. Day, Heather J. Cordell, Baki Topal, Karine Clément, Stefano Romeo, Vlad Ratziu, Tania Roskams, Ann K. Daly, Quentin M. Anstee, Matthias Trost, Anetta Härtlova

**Affiliations:** Translational and Clinical Research Institute, Faculty of Medical Sciences, Newcastle University, Newcastle upon Tyne, United Kingdom; Wallenberg Centre for Molecular and Translational Medicine, Department of Microbiology and Immunology at Institute of Biomedicine, University of Gothenburg, Gothenburg, Sweden; Biosciences Institute, Faculty of Medical Sciences, Newcastle University, Newcastle upon Tyne, United Kingdom; Radiation Oncology, Albert Einstein College of Medicine, New York, USA; Center for Brain & Disease Research, VIB-KU Leuven, Leuven, Belgium; Department of Human Genetics, KU Leuven, Leuven, Belgium; Department of Imaging and Pathology, Translational Cell and Tissue Research, KU Leuven and University Hospitals Leuven, Leuven, Belgium; The Wallenberg Laboratory for Cardiovascular and Metabolic Research, Department of Molecular and Clinical Medicine, University of Gothenburg, Gothenburg, Sweden; Nutrition and obesity: systemic approaches, Inserm, Sorbonne University, Paris, France; Population Health Sciences Institute, Faculty of Medical Sciences, Newcastle University, Newcastle upon Tyne, United Kingdom; Department of Medical Sciences, Division of Gastro-Hepatology, A.O. Città della Salute e della Scienza di Torino, University of Turin, Turin, Italy; Department of Pathology, Aretaieio Hospital, National & Kapodistrian University of Athens, Athens, Greece; NIHR Nottingham Biomedical Research Centre, Nottingham University Hospitals NHS Trust and University of Nottingham, Nottingham, United Kingdom; Liver Unit, Department of Medicine, Cambridge Biomedical Research Centre, Cambridge University NHS Foundation Trust, United Kingdom; University of Cambridge Metabolic Research Laboratories, Wellcome-MRC Institute of Metabolic Science, Addenbrooke’s Hospital, Cambridge, United Kingdom; Bioscience COPD/IPF, Research and Early Development, Respiratory and Immunology (R&I), BioPharmaceuticals R&D, AstraZeneca, Gothenburg, Sweden; Department of Medicine, University of Helsinki, Helsinki, Finland; University Clinic for Visceral Surgery and Medicine, University of Bern, Bern, Switzerland; Hepatology, Department of Biomedical Research, University of Bern, Bern, Switzerland; Division of Gastroenterology and Hepatology, Department of Medicine and Health Sciences, Linköping University, Linkoping, Sweden; Department of Gastroenterology and Hepatology, Antwerp University Hospital & University of Antwerp, Antwerp, Belgium; Sezione di Gastroenterologia, Dipartimento Biomedico di Medicina Interna e Specialistica, Università di Palermo, Palermo, Italy; I. Department of Medicine, University Hospital Mainz, Mainz, Germany; Department of Abdominal Surgery, KU Leuven and University Hospitals Leuven, Leuven, Belgium; Assistance Publique-Hôpitaux de Paris, hôpital Beaujon, University Paris-Diderot, Paris, France; Newcastle NIHR Biomedical Research Centre, Newcastle upon Tyne Hospitals NHS Trust, Newcastle upon Tyne, United Kingdom

## Abstract

Obesity-associated inflammation is a key player in the pathogenesis of non-alcoholic fatty liver disease (NAFLD). However, the exact mechanisms remain incompletely understood. Here we demonstrate that macrophage scavenger receptor 1 (MSR1, CD204) expression is associated with the occurrence of hepatic lipid-laden foamy macrophages and correlates with the degree of steatosis and steatohepatitis in a cohort of 170 NAFLD patients. Mice lacking Msr1 are protected against high fat-cholesterol diet (HFD)-induced metabolic disorder, showing fewer hepatic lipid-laden foamy macrophages, less hepatic inflammation, improved dyslipidemia and glucose tolerance, while showing a change in hepatic lipid metabolism. We show that MSR1 induces a pro-inflammatory response via the JNK signaling pathway upon triggering by saturated fatty acids. *In vitro* blockade of the receptor prevented the accumulation of lipids in primary macrophages which inhibited the switch towards a pro-inflammatory phenotype and the release of cytokines such as TNFa. Targeting MSR1 using monoclonal antibody therapy in an obesity-associated NAFLD mouse model and *ex vivo* human liver slices resulted in the prevention of foamy macrophage formation and liver inflammation. Moreover, we identified that rs41505344, a polymorphism in the upstream transcriptional region of *MSR1,* was associated with altered serum triglycerides and aspartate transaminase levels in a cohort of over 400,000 patients. Taken together, our data suggest a critical role for MSR1 in lipid homeostasis and a potential therapeutic target for the treatment of NAFLD.

**One Sentence Summary:** The immunometabolic role of MSR1 in human NAFLD.

## Introduction

With the increasing prevalence of obesity, non-alcoholic fatty liver disease (NAFLD) has become the most common chronic liver disease globally (*1*). NAFLD is characterized by excessive hepatic triglyceride accumulation and represents a series of diseased states ranging from isolated steatosis (non-alcoholic fatty liver, NAFL) to non-alcoholic steatohepatitis (NASH), identified by the presence of necro-inflammation and hepatocyte ballooning, with varying degrees of fibrosis. NAFLD is strongly linked with metabolic syndrome, i.e. dyslipidemia, hypertension, obesity and type 2 diabetes mellitus (T2DM), and currently affects 20 to 30% of the global population (*2*). Importantly, not all patients progress from NAFL to NASH and although gene signatures of more advanced fibrosing-steatohepatitis have been identified, the exact pathogenic pathways involved in the initiating phases of the disease, especially the transition from NAFL to NASH, are not fully understood (*3*).

Growing evidence supports the view that Kupffer cells, the endogenous hepatic macrophages, are initiators of inflammation and hence contribute to NAFLD development, whilst recruited monocyte-derived macrophages play a crucial role in the disease progression (*4, 5*). Hepatic macrophages are responsive to a variety of stimuli including bacterial endotoxins (such as lipopolysaccharide) but also free fatty acids (FFAs) or cholesterol (*6*). Excess of FFAs and cholesterol can cause the formation of hepatic foamy macrophages, resembling those seen in atherosclerotic lesions, and leads to Kupffer cell aggregates and lipogranulomas during steatohepatitis (*7, 8*). Specifically the intake of saturated fat has been shown to induce insulin resistance, enhance intrahepatic triglyceride accumulation and steatohepatitis (*9*). Liver biopsies from patients with metabolic NAFLD have been described to be markedly enriched in saturated and monounsaturated triacylglycerols and FFAs (*10*). In advanced NASH, specific individual saturated FAs (SFAs), such as myristic acid (C14:0) or palmitic acid (PA, C16:0), proved to be increased when compared to earlier stages of the disease (*11*). However, the molecular mechanisms underlying hepatic macrophage activation and/or the formation of lipid-laden foamy macrophages in NAFLD remain poorly understood.

PA, rather than non-SFAs, has been shown to be a strong inducer of inflammation in immortalized cell lines through activation of the downstream JNK signaling pathway (*12, 13*). Recent data show that pro-inflammatory activation of murine bone marrow-derived macrophages by PA is independent of Toll-like receptor 4, yet the receptor that is responsible is still not known (*14*). Recently, we have shown that *in vitro* activation of the phagocytic receptor, macrophage scavenger receptor 1 (MSR1, also known as SR-A or CD204), results in pro-inflammatory macrophage polarization through JNK activation (*15*). MSR1 is a key macrophage receptor for the clearance of circulating lipoproteins and has been implicated in atherogenesis (*16, 17*). We therefore hypothesized that MSR1 might be involved in inflammatory responses in the context of lipid overload during obesity-induced NAFLD.

Here we show that *MSR1* gene expression is associated with NAFLD disease activity in a large cohort of patients with histologically proven NAFLD and that MSR1 deficiency in a mouse model of obesity-associated NAFLD prevents foamy macrophage formation, and the induction of liver inflammation and fibrosis. Moreover, we found that systemic administration of MSR1 monoclonal antibody could modify NAFLD pathology in a mouse model and human *ex vivo* liver slices of obesity-associated NAFLD.

## Results

### MSR1 expression correlates with steatohepatitis activity in human NAFLD

As MSR1 is a receptor for lipids in macrophages, we investigated the impact of MSR1 on NAFLD. We first analyzed *MSR1* gene expression in a cohort of 170 histologically characterized human adult liver biopsies using nanoString transcriptomics. The cohort was stratified according to histopathological disease grade and stage, i.e. NAFL and NASH with fibrosis ranging from F0 to F4 (**Table S1**). The analysis revealed a significant increase of *MSR1* gene expression in NASH compared to NAFL (p<0.01), indicating a positive correlation with the active inflammatory form of the disease (**Fig. 1a**). This observation was supported by a significant correlation between *MSR1* mRNA expression and steatosis grade (Spearman r=0.29, p=0.0001), hepatocyte ballooning (r=0.22, p<0.01) and lobular inflammation (r=0.24, p<0.01), while there was no correlation with stages of hepatic fibrosis indicating that *MSR1* mRNA are strongly linked to steatohepatitis activity rather than wound healing *per se* (**Fig. 1b**). In addition, *MSR1* mRNA correlated with the NASH Clinical Research Network NAFLD Activity Score (NAS), defined as the sum of steatosis, ballooning and lobular inflammation (r=0.35, p<0.0001, **Fig. S1a**) (*18*). Patients with NAS≥4 showed a significant increase in MSR1 expression (fold change =1.34, p<0.0001, **Fig. S1a**). Furthermore, *MSR1* positively correlated with serum levels of alanine transaminase (ALT, r=0.31, p=0.0001) and aspartate transaminase (AST, r=0.32, p<0.0001) (**Fig. S1b**). Histopathological analysis of human adult NAFLD liver biopsies showed that MSR1 was predominantly expressed in resident liver macrophages, the Kupffer cells, rather than infiltrating monocyte-derived macrophages located in the portal tract, as visualized by the MSR1 and CD68 immunostaining (**Fig. 1c and Fig. S2a-b**). While the number of infiltrating portal CD68-immunopositive cells increased with disease progression (p<0.05), no significant differences were found for infiltrating MSR1-positive cells (**Fig.1c**). These results were confirmed using publicly available single cell RNA sequencing data from healthy and end-stage cirrhotic liver, showing that *MSR1* expression was mainly restricted to the Kupffer cell population, whereas *CD68* or the expression of other scavenger receptors, such as *CD36*, was also found in monocyte populations (**Fig. S2c**) (*19*). Notably, MSR1 immunopositivity was seen in lipid-laden foamy macrophages and lipogranulomas throughout the spectrum of NAFLD, indicating a role for MSR1 in lipid accumulation in endogenous macrophages and concordantly inflammation (**Fig. 1c and Fig. S2d**).

**Fig. 1.**
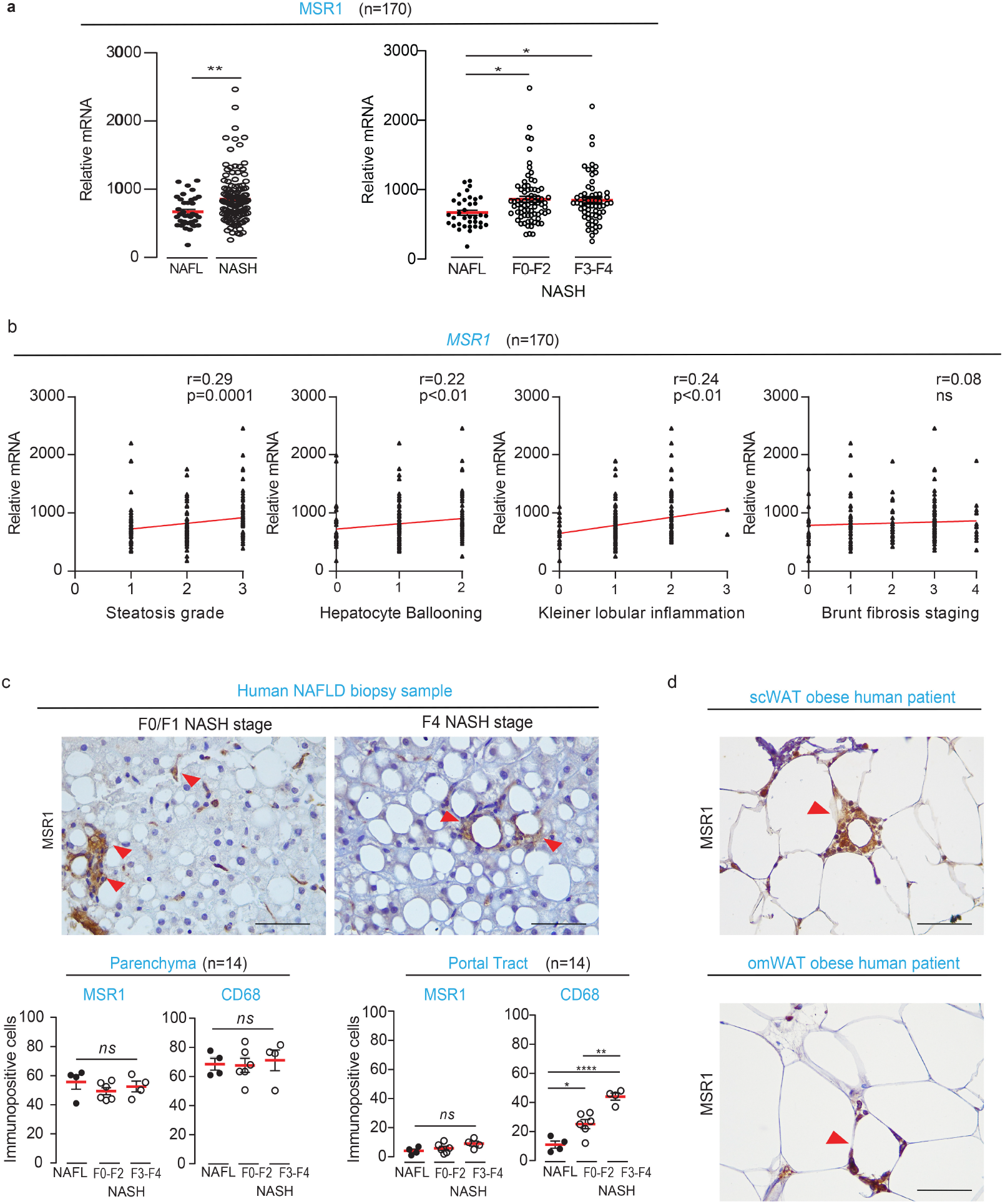
Macrophage Scavenger Receptor 1 (MSR1) expression in human non-alcoholic fatty liver disease (NAFLD) correlates with steatosis and steatohepatitis. (**a**) mRNA levels of *MSR1* in a cohort of 170 histological proven NAFLD samples covering the different stages of the disease (NAFL non-alcoholic fatty liver; NASH non-alcoholic steatohepatitis F0-4 fibrosis stage) using nanoString. Filled symbols, NAFL stage; open symbols, NASH stage. (Mann-Whitney-U test and Kruskal-Wallis with correction for multiple testing) (**b**) Correlation of *MSR1* gene expression with the steatosis, hepatocyte ballooning, lobular inflammation and fibrosis stage (Spearman correlation) (**c**) The immunohistochemical analysis of MSR1 in liver tissue of human NAFLD biopsies (n=14), red arrows indicate foamy macrophages and lipogranuloma. Histopathological quantification of MSR1 and CD68 immunopositive cells in the parenchyma and portal tract of human NAFLD samples with mild and advanced fibrosis (NAFL n=4; NASH F0-2 n=6; NASH F3-4 n=4; one-way ANOVA or Kruskal-Wallis with correction for multiple testing based on distribution). (**d**) Representative image of immunohistochemical staining for MSR1 in subcutaneous and omental white adipose tissue (scWAT/omWAT) from an obese patient. Data are presented as mean ± SEM (* p<0.05, ** p<0.01, *** p<0.001, **** p<0.0001, *ns:* non-significant). Scale bars 100μm.

As macrophages have been reported to regulate metabolic homeostasis in adipose tissue (*20*), we evaluated MSR1 expression in subcutaneous and matched omental adipose tissue from obese patients (BMI >30 kg/m^2^). MSR1-immunopositive macrophages were observed in crown-like structures in both types of adipose tissue, though no association was found between MSR1-immunopositive cells present in the adipose tissue and the diagnosis of NASH (**Fig. 1d and Fig. S2e**). Taken together, these human data demonstrate a positive correlation of *MSR1* transcript and protein levels with obesity-associated NAFLD and occurrence of hepatic-resident lipid-laden macrophages in the presence of excess lipid accumulation.

### Msr1 deficiency protects against diet-induced metabolic dysregulation and liver damage in mice

To further investigate how MSR1 functionally contributes to the development of obesity-related NAFLD, we subjected *Msr1*^-/-^ mice (n=5) and their corresponding *Msr1*^+/+^ (n=5, wild-type, WT) male and age-matched counterparts to a high-fat and high-cholesterol diet (HFD) for 16 weeks. Upon HFD feeding, *Msr1*-deficient mice displayed an increased total body weight and an increase in liver and epididymal white adipose tissue (eWAT) weight compared to WT (p<0.05, **Fig. 2a-b**). Consistent with enhanced adipose mass, HFD-fed *Msr1*^-/-^ mice displayed enhanced fatty acid accumulation within the cells (p<0.0001, **Fig. 2c**) along with larger adipocytes compared to WT mice (p<0.0001, **Fig. 2d-e**), indicating an increased adiposity and fat storage in the absence of *Msr1*. Although no murine models accurately recapitulate all histological features of human steatohepatitis (*21*), histological and transcriptomic features of liver fibrosis were clearly attenuated by *Msr1* deficiency upon HFD feeding (**Fig. 2d-f**). Sixteen weeks of regular diet did not result in any histological differences between the livers of WT and *Msr1*^-/-^ mice (**Fig. S3a**), while WT mice on HFD displayed a significant higher hepatic fibrosis stage (p<0.05, **Fig. S3a-b**) and increased collagen deposition (p<0.01, **Fig. 2e**) compared to the *Msr1*^-/-^ mice. Next, we characterized the livers of HFD-WT and HFD-*Msr1*^-/-^ mice by high-throughput RNA sequencing analysis. The analysis revealed 728 differentially expressed genes (adjusted p-value<0.05, **Fig. S3c**, **Table S2**). Gene Ontology analysis of differentially expressed genes highlighted an enrichment for genes correlating to biological processes including “immune system process”, “innate immune response”, “phagocytosis” and “lipid metabolic process” (**Fig. 2f**, **Fig.S3d**). *Msr1*^-/-^ mice displayed a reduced hepatic transcript expression of inflammatory cytokines (including *Axl, Ccl6, Il1b, Spp1*), pro-inflammatory immune cell markers (*Ccr5, Cd14, Cd44, S100a8, S100a9*), markers for hepatic stellate cell activation (*Sox9, Pdgfb*) and members of the *Tnfa* signaling pathway (*Ripk3, Tnfaip2, Tnfaip8l2*) when compared with WT mice after 16 weeks of HFD (**Fig.2f**). Furthermore, *Msr1*^-/-^ mice on HFD showed a shift in gene expression associated with lipid metabolism, with genes including *Acox1, Acox2, Apoe, Ces1d, Hsd17b11, Pla2g6* and *Ppara* increasing, and genes such as *Fabp5, Lpcat2, Lpl, Pla2g7* and *Pnpla3* decreasing (**Fig.2f**). To further test the effect of *Msr1-deficiency* on the metabolic syndrome, we measured several parameters within the serum as well as in the liver tissue. HFD-fed *Msr1*^-/-^ mice exhibited higher serum leptin (p<0.05), lower concentrations of circulating FFAs (p<0.05) which corresponded to reduced hepatic triglycerides (TGs, p<0.01) and enhanced lipid utilization measured by approximately 50% higher mitochondrial oxygen consumption rate in the liver compared to HFD-fed WT mice (p<0.001) (*22*), while circulating TG levels remained unchanged (**Fig. 2g-h, Fig. S3e**). Moreover, the absence of *Msr1* improved glucose uptake from blood in obese mice (p<0.01, **Fig. S3f**). Taken together, these results demonstrate that *Msr1* deficiency increases the body weight but protects against features of the metabolic syndrome, including liver inflammation and fibrosis, while modulating hepatic lipid metabolism.

**Fig. 2.**
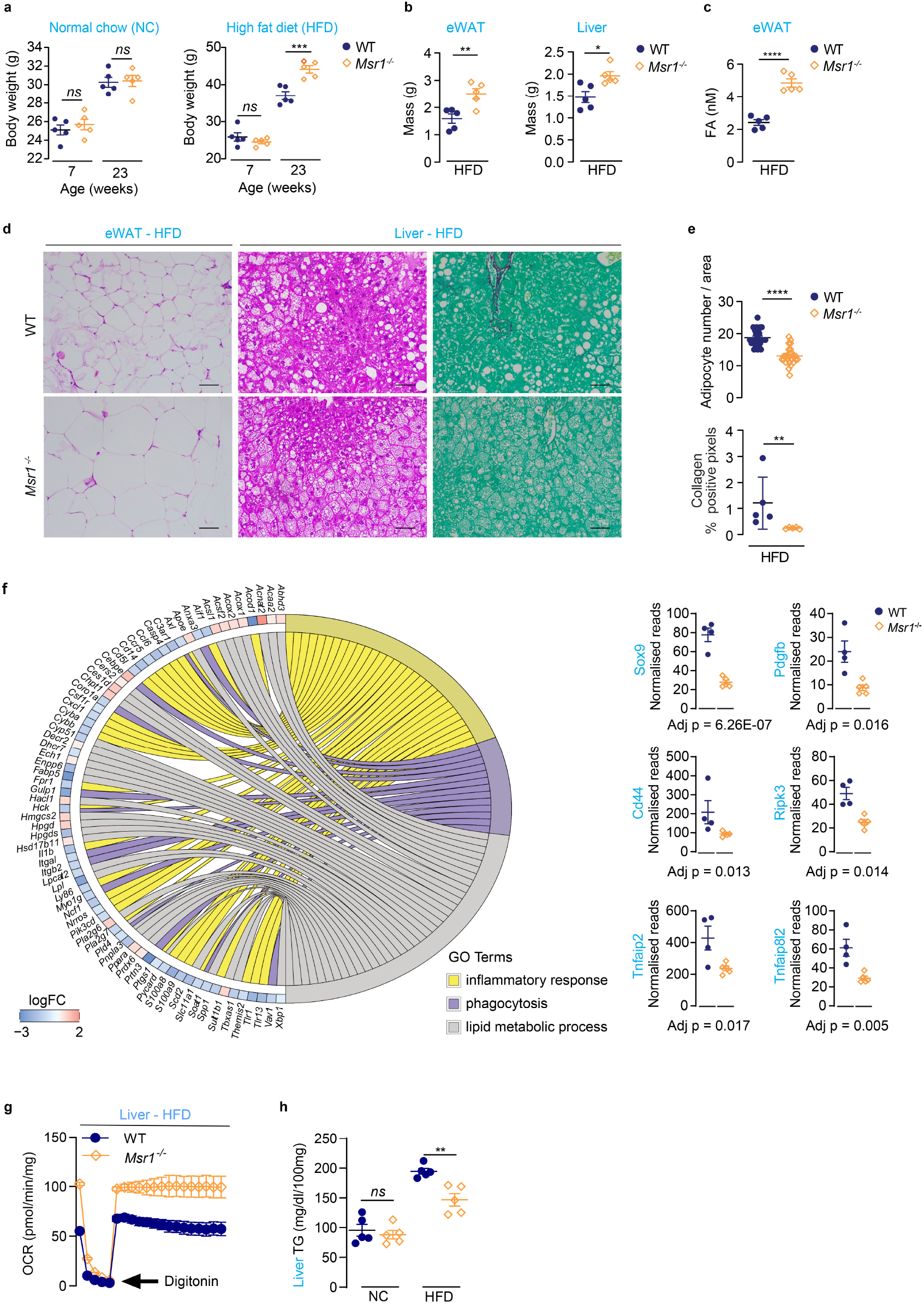
Msr1 deficiency protects against HFD-associated metabolic dysregulation and liver damage. (**a**) Body weight of *Msr1*+/+ (Wild-type, WT) and *Msr1*-/-male aged-matched mice fed either with normal control chow diet (NC) or high fat, high cholesterol diet (HFD) for 16 weeks (n=5 mice/experimental group) (one-way ANOVA with correction for multiple testing; p-values are shown for the comparisons WT and *Msr1*-/-). (**b**) Epididymal (eWAT) and liver mass of WT and *Msr1*-/-male mice fed with HFD (unpaired Student’s t-test). (**c**) Quantification of fatty acids in eWAT of HFD fed mice (unpaired Student’s t-test). (**d**) Representative images of morphology of the eWAT and liver from HFD-fed WT and *Msr1*-/- mice, as assessed by H&E staining and Picro Sirius Red Fast Green staining. Scale bar 100μm. (**e**) Quantification of the adipocyte number per area and hepatic collagen deposition of WT and *Msr1*^-/-^ HFD-fed mice (n = 5 mice/experimental group; Mann-Whitney-U test). (**f**) RNA sequencing data comparing Msr1-/- (n=5) with baseline WT (n=4) HFD-fed mice. Gene Ontology enrichment analysis was performed for biological processes and selected differentially expressed genes were visualized. (**g**) Seahorse analysis of oxygen consumption rates (OCRs) and area under curve (AUC) of HFD-WT and HFD-fed *Msr1*-/- aged-matched male mice (n = 4/experimental group). (**h**) Quantification of triglycerides (TG) in liver tissue samples (n = 5/experimental group; one-way ANOVA with correction for multiple testing; p-values are shown for the comparisons WT and *Msr1*-/-). Each symbol represent animal. Filled symbols, Msr1+/+ (WT); open symbols, *Msr1*-/-mice. Data are presented as mean ± SEM (*p< 0.05, **p < 0.01, ***p < 0.001, ****p < 0.0001, ns: non-significant).

### Msr1 deficiency prevents formation of pro-inflammatory foamy macrophages in vivo

Next, we asked whether the lipid-laden environment is a proximal stimulus leading to Msr1-mediated inflammation in the liver and adipose tissue, which may explain the observed metabolic dysfunction. In agreement with our human data, histopathological analysis of the liver and adipose tissue from HFD-fed *Msr1*^-/-^ mice showed no hepatic lipogranuloma and very few foamy macrophages compared to their WT counterparts, demonstrated by F4/80 immunostaining (**Fig. 3a**). We have recently shown that triggering of Msr1 activates the JNK signaling pathway (*15*). JNK signaling is a stress-activated pathway that has been implicated in SFA-induced pro-inflammatory activation of macrophages (*14*). Indeed, HFD-fed *Msr1*^-/-^ mice exhibited reduced hepatic JNK1/2 phosphorylation compared to the WT (**Fig. S4a**). Moreover, *Msr1*^-/-^ mice displayed lower Il6 and Tnfa serum levels and reduced *Tnfa* and *Il6* gene expression in the liver and eWAT (**Fig. 3b-d**). Furthermore, *Msr1* deficiency impaired pro-inflammatory activation of isolated adipose tissue-(ATMs) and hepatic-associated macrophages as shown by lower gene transcripts of *Tnfa* and *Il6* (p<0.01, **Fig. 3e-g**). To extend these findings, we co-cultured either primary adipocytes and hepatocytes or differentiated 3T3-L1 adipocyte-like cells and lipid-fed hepatocyte-like Hepa1-6 cells indirectly with WT and *Msr1*^-/-^ bone marrow-derived macrophages in trans-well assays (**Fig. S4b-d**). Consistent with the *in vivo* data, *Msr1* deficiency or blocking of the Msr1 receptor using a monoclonal antibody attenuated the pro-inflammatory response when co-cultured with adipocytes or hepatocytes compared to WT counterpart (**Fig. S4d-f**). Altogether, these results show that Msr1 mediates HFD-induced hepatic and adipose tissue inflammation and facilitates macrophage activation toward a pro-inflammatory phenotype.

**Fig. 3.**
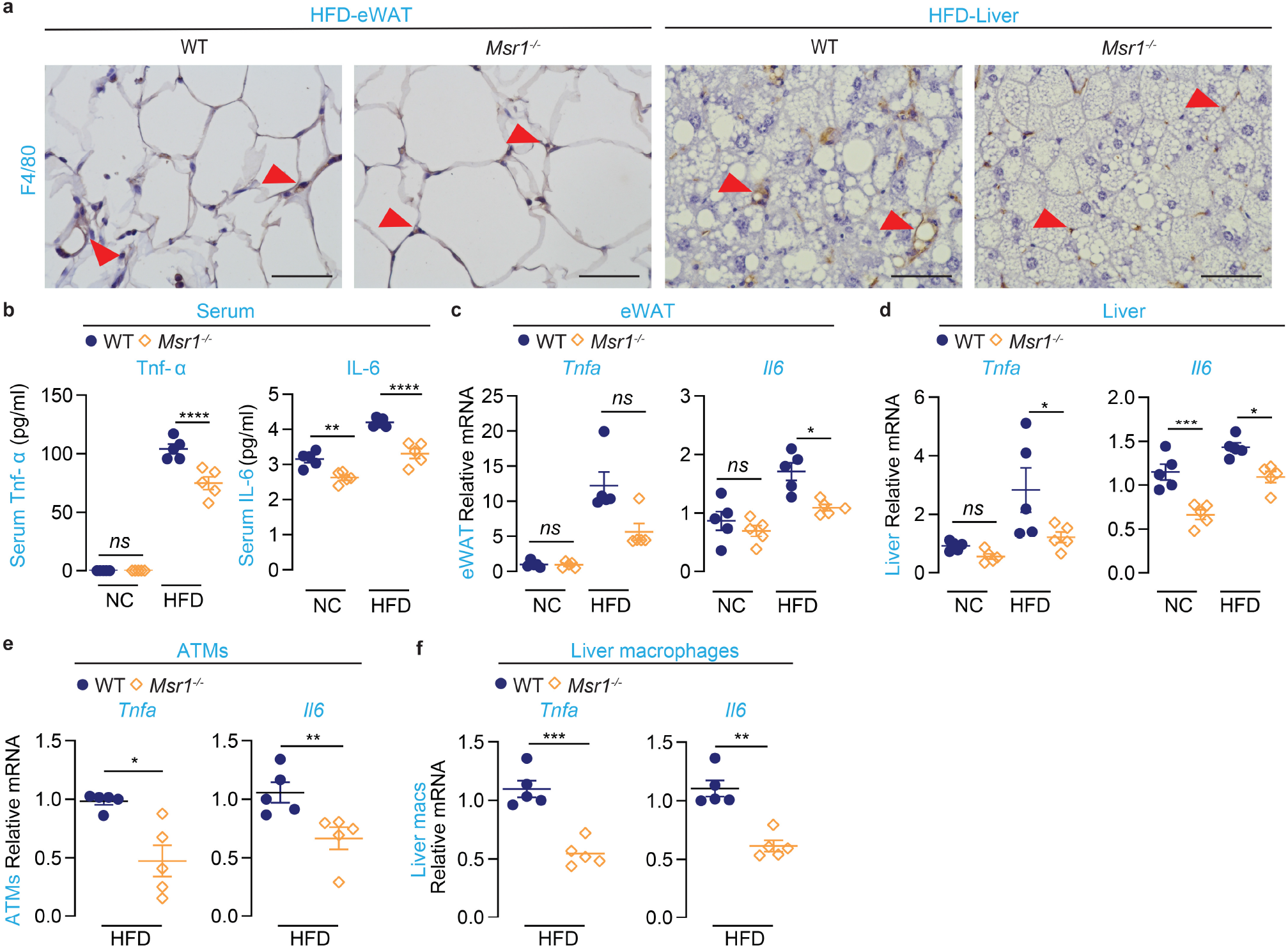
Msr1 mediates HFD-induced adipose tissue and hepatic inflammation and facilitates macrophage activation toward a pro-inflammatory phenotype. (**a**) Representative images for F4/80 immunostainings in eWAT and liver tissue (scale bars 100μm) from WT and *Msr1*^-/-^ HFD-fed male aged-matched mice (n=5/experimental group). Arrows indicate immunopositive cells. (**b**) Serum levels of Tnfa and Il-6 in NC-and HFD-fed mice (n=5/experimental group). (**c-d**) Quantification of mRNA levels of *Tnfa, Il6* inflammation markers in the eWAT and liver of NC-and HFD-fed mice (n=5/experimental group). (**e-f**) Real-time PCR analysis for markers of inflammation in F4/80^+^ adipose tissue (ATMs) and liver macrophages from WT and *Msr1*^-/-^ HFD-fed aged-matched male mice (n=5 mice/experimental group). Each symbol represent animal. Filled symbols, (WT); open symbols, *Msr1*^-/-^ mice. Data are presented as mean ± SEM (unpaired Student’s t-test or Mann-Whitney-U test, or one-way ANOVA or Kruskal-Wallis with correction for multiple testing based on the distribution of the data; p-values are shown for the comparisons WT and *Msr1*-/-; *p< 0.05, **p < 0.01, ***p < 0.001, ****p < 0.0001, ns: non-significant).

### Triggering of Msr1 by lipids induces JNK-mediated pro-inflammatory activation of macrophages

We next investigated the underlying mechanism of Msr1-mediated lipid-induced inflammation. We reasoned that Msr1 is directly responsible for lipid uptake in macrophages, leading to an inflammatory response independent from other cell types. In this regard, we measured the uptake of SFA (palmitic acid) and non-SFA (oleic acid) in either hepatic macrophages or bone marrow-derived macrophages (BMDMs) by quantifying Oil-red-O staining using confocal microscopy (**Fig. 4a-c**). The analysis revealed that Msr1 facilitates the uptake of both SFA as well as non-SFA but only SFA induced enhanced levels of *Tnfa* and *Il6* transcripts and phosphorylation of JNK signaling in WT macrophages (**Fig. 4d,e, Fig. S5a-c**). Furthermore, blocking of Msr1 receptor with a monoclonal antibody or the chemical inhibition of JNK signaling in macrophages abrogated the induction of *Tnfa* and *Il6* pro-inflammatory gene expression in response to SFA treatment (**Fig. 4f**, **Fig.S5d**). These data indicate that SFA-induced triggering of Msr1 regulates JNK-mediated pro-inflammatory activation of macrophages in the absence of lipopolysaccharide.

**Fig. 4.**
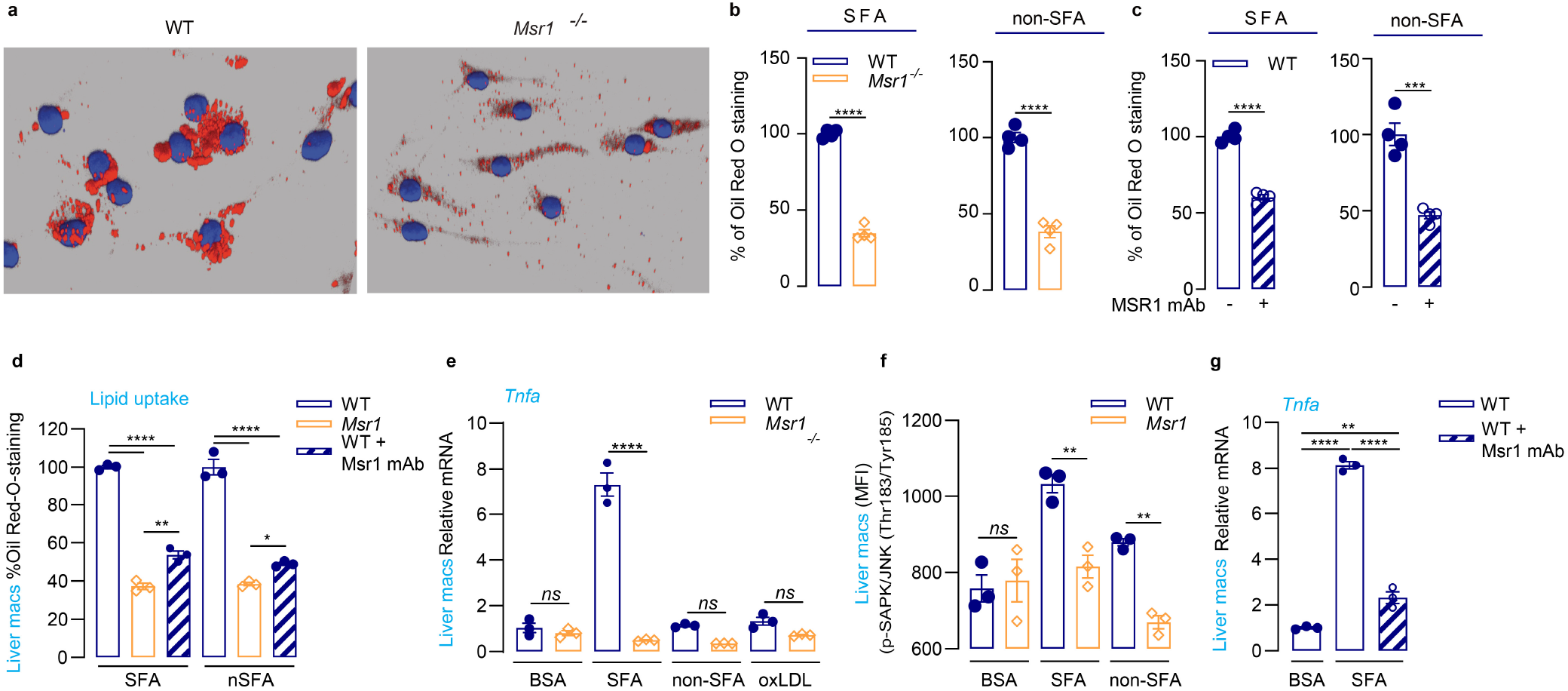
Msr1 regulates JNK-mediated lipid-induced pro-inflammatory activation of macrophages. (**a**) Representative image of lipid uptake (mixture of 1mM Palmitic acid, saturated fatty acids (SFA), and 2mM Oleic, non-saturated fatty acids (non-SFA)) by WT and *Msr1*^-/-^ bone marrow derived macrophages (BMDMs) visualized by Oil-red-O staining using confocal microscopy (n=3). (**b-c**) Quantification of SFA (palmitic acid 1mM) and non-SFA (oleic acid 2mM) uptake either by WT and *Msr1*^-/-^ BMDMs, or WT BMDMs pre-treated with or without monoclonal anti-Msr1 antibody using Oil-red-O staining (n=4). Data are normalized to the average of the WT BMDM group. (**d**) Quantification of Oil-red-O staining to assess the uptake of SFA (palmitic acid 1mM) and non-SFA (oleic acid 2mM) by WT and *Msr1*^-/-^ primary liver macrophages, or WT primary liver macrophages pre-treated with or without monoclonal anti-Msr1 antibody (n=4). Data are normalized to the average of the WT BMDM group. (**e**) Real time-PCR analysis of *Tnfa* pro-inflammatory gene in WT and *Msr1*^-/-^ primary liver macrophages treated either with control bovine serum albumin (BSA) or SFA, non-SFA or oxidized low-density lipoprotein (oxLDL) for 6hrs (n=3). (**f**) Flow cytometry analysis of phospho-JNK (Thr183/Tyr185) in WT and *Msr1*^-/-^ primary liver macrophages treated either with BSA as control or SFA and non-SFA for 3hrs (n=3). (**g**) Real time-PCR analysis of *Tnfa* expression in WT primary liver macrophages pre-treated with or without monoclonal anti-Msr1 antibody (n=3). Data are shown as mean ± SEM (unpaired Student’s t-test or one-way ANOVA with correction for multiple testing; for panel d-f, the p-values are shown only for grouped comparisons per experimental condition; *p< 0.05, **p < 0.01, ***p < 0.001, ****p < 0.0001).

### Therapeutic inhibition of MSR1 prevents formation of pro-inflammatory foamy macrophages

To investigate the therapeutic potential of targeting MSR1 in the treatment of NAFLD, we applied an antibody-based intervention using NAFLD mouse models and *ex vivo* human liver slices. WT mice were fed a HFD for 12 weeks and were administered two doses of monoclonal rat anti-mouse Msr1 antibody (n=3 animals) or isotype-matched IgG control (n=4 animals) at week 10 and 11 by intravenous injection (**Fig. 5a**). Histological assessment by F4/80 staining showed a reduction in foamy macrophages and lipogranulomas in the livers of mice treated with anti-Msr1compared to the IgG control (**Fig. 5b**). Digital quantification of the F4/80 stainings indicated a significant reduction in surface area positivity of hepatic macrophages in the anti-Msr1 treated animals (p<0.05, **Fig.5c**). Furthermore, treated animals showed reduced expression of *Tnfa* transcript in liver samples and isolated hepatic macrophages (p<0.05, **Fig.5d**).

**Figure 5.**
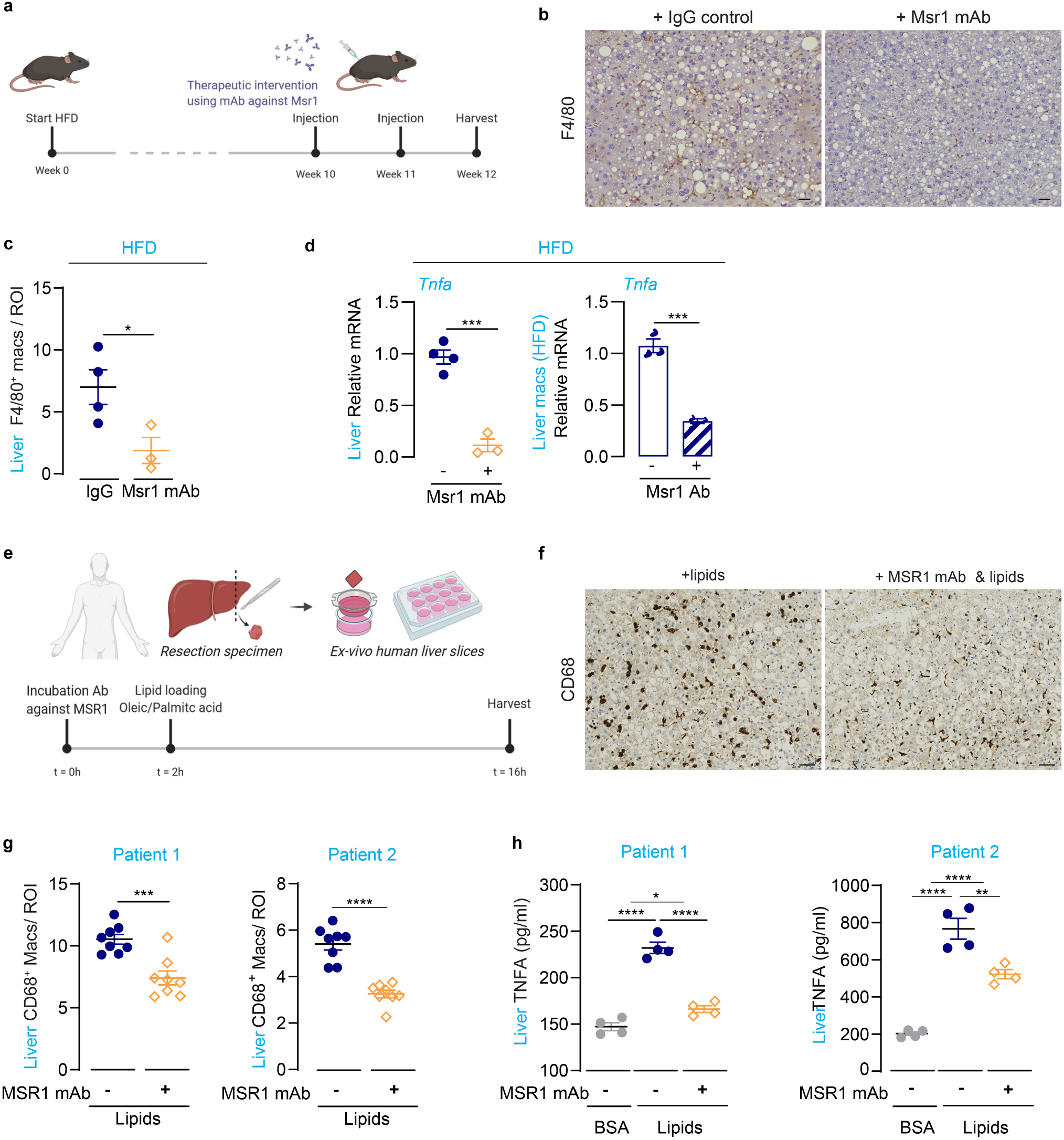
Therapeutic inhibition of MSR1 prevents formation of pro-inflammatory foamy macrophages. Schematic overview of therapeutic antibody-based intervention in a NAFLD mouse model. (**b**) Immunohistochemical staining of liver samples for F4/80 from HFD-fed animals treated with anti-Msr1 antibody (n=3) or IgG control (n=4). (**c**) Quantification of F4/80 staining from treated animals presented as percentage pixel positivity of the region of interest (ROI). (**d**) Real-time PCR analysis for *Tnfa* transcript in liver samples and isolated hepatic macrophages from HFD-fed animals treated with anti-Msr1 antibody (n=3) or IgG control (n=4). Isolated primary liver macrophages were pooled together before real-time PCR analysis (n=3). (**e**) Schematic overview of antibody-based treatment of *ex vivo* lipid-loaded human liver slices. Per patient, two biological replicates were used for each condition. Samples were loaded with a combination of oleic (2mM) and palmitic acid (1mM). (**f**) Immunohistochemical staining for CD68 on human lipid-loaded liver slices treated with or without anti-MSR1 antibody. (**g**) Quantification of CD68 staining (4 areas per liver slice) presented as percentage pixel positivity of the region of interest (ROI). (h) TNFa ELISA from human liver slices treated with control Bovine Serum Albumin (BSA), lipids or anti-MSR1-antibody+lipids (two technical measurements were performed per biological replicate). Data are presented as mean ± SEM (unpaired Student’s t-test or one-way ANOVA with correction for multiple testing; *p< 0.05, **p < 0.01, ***p < 0.001, ****p < 0.0001). Scale bars 50μm.

To further investigate whether inhibition of MSR1 prevents the formation of foamy macrophages in humans, we collected human liver slices with normal morphology from two different patients (2 biological replicates per condition for each patient sample). The samples were incubated with a polyclonal rabbit anti-human MSR1 antibody prior to culturing them with a mixture of OA (2mM) and PA (1mM) in combination with anti-MSR1 antibody for a total duration of 16h (**Fig.5e**). Treatment with the antibody reduced the accumulation of lipids within the macrophages as shown by the CD68 immunohistochemical staining (**Fig.5f**). Digital quantification of the CD68 staining showed a significant reduction in surface area positivity of Kupffer cells upon antibody treatment in both patient samples as compared to the non-treated condition (p<0.001, four areas quantified per biological replicate, **Fig.5g**). Moreover, lipid loading induced the release of TNFa into the cell culture medium as compared to the bovine serum albumin (BSA) control in both patient samples (p<0.0001, two technical replicates per biological replicate), which was significantly reduced upon treatment with anti-MSR1 antibody (p<0.01)(**Fig.5h**). Overall, our *in vivo* and *ex vivo* results show that therapeutic inhibition of MSR1 prevents the formation of foamy macrophages and the release of TNFa.

### Relevance of polymorphisms in *MSR1* region to NAFLD and metabolic traits

Next, we asked whether genetic variants in MSR1 are associated with susceptibility to NAFLD and if there is an association with transcriptional regulatory mechanisms controlling *MSR1* expression. Using previously published genomics data encompassing a cohort of 1,483 European Caucasian patients with histologically proven NAFLD and 17,781 European general-population controls (*23*), we identified 4 single nucleotide polymorphisms (SNPs) in or around the *MSR1* locus with p-values<5*10^-4^, with rs41505344 as the most significant (p=1.64*10^-4^) (**Fig. 6a** and **Table S3**). Quantitative trait analysis for rs41505344 in 430,101 patients enrolled in the UKBiobank showed a significant correlation with serum triglycerides and aspartate transaminase (AST) levels, even after adjustment for age, gender, body mass index (BMI), center, batch and the first ten principal components (**Table 1**, adjusted p=3.55*10^-6^ and adjusted p=0.003 respectively).

**Figure 6.**
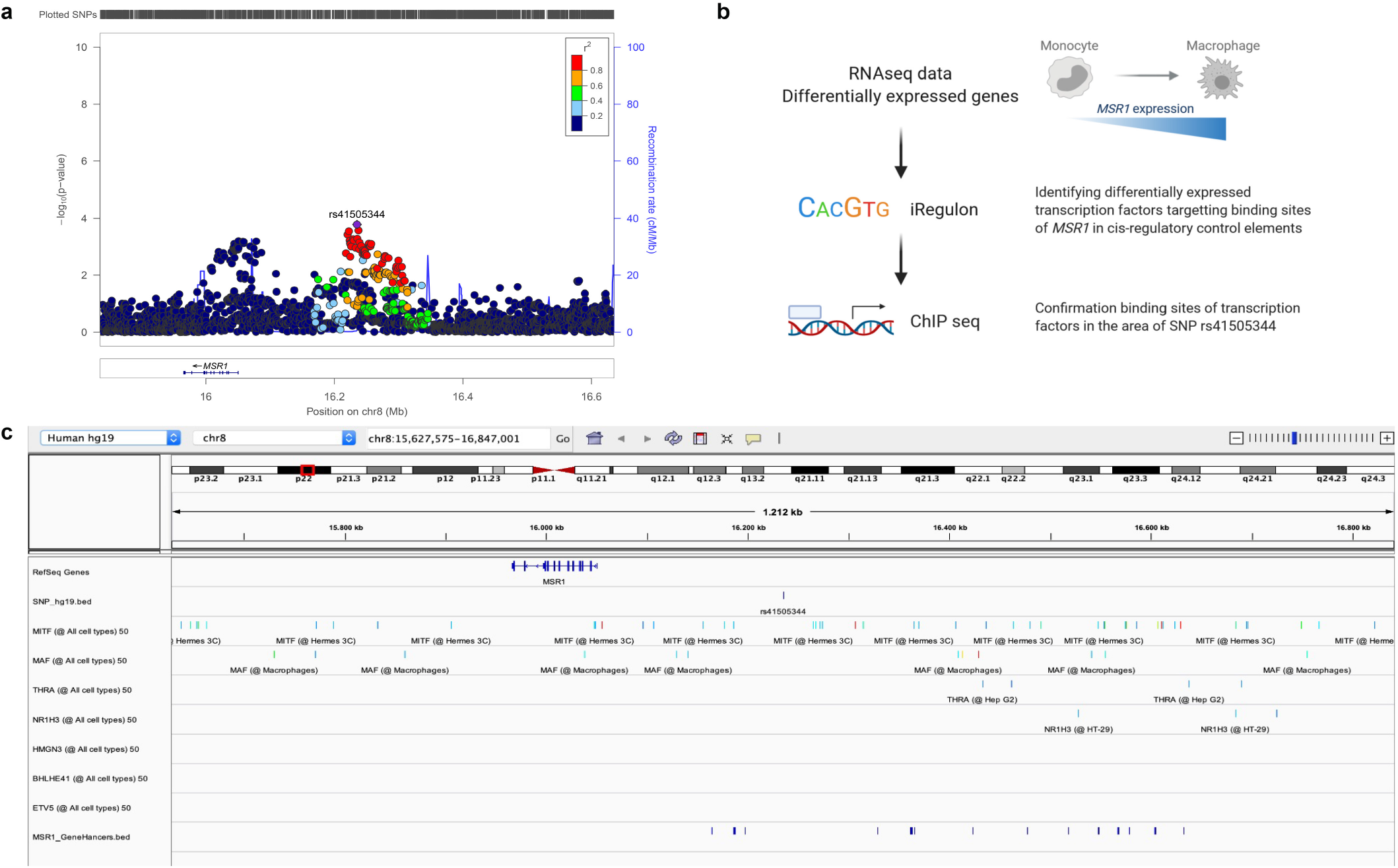
Regulatory mechanisms of *MSR1* expression in human NAFLD. (**a**) Locus plot showing *MSR1* rs41505344 SNP based on case-control analysis comparing 1,483 histologically characterized NAFLD samples with 17,781 matched population controls. (**b**) Schematic overview of the workflow used to identify transcriptional regulatory mechanisms of *MSR1* from publicly available RNA sequencing data, comparing human monocytes with differentiated macrophages (*24*). (**c**) Visualization of chromatin immunoprecipitation sequencing data around *MSR1* rs41505344 SNP of the predicted transcription factors that are differentially expressed in the RNA sequencing data as identified by iRegulon. Bottom row indicates known transcriptional regulatory regions of *MSR1*.

**Table 1.**
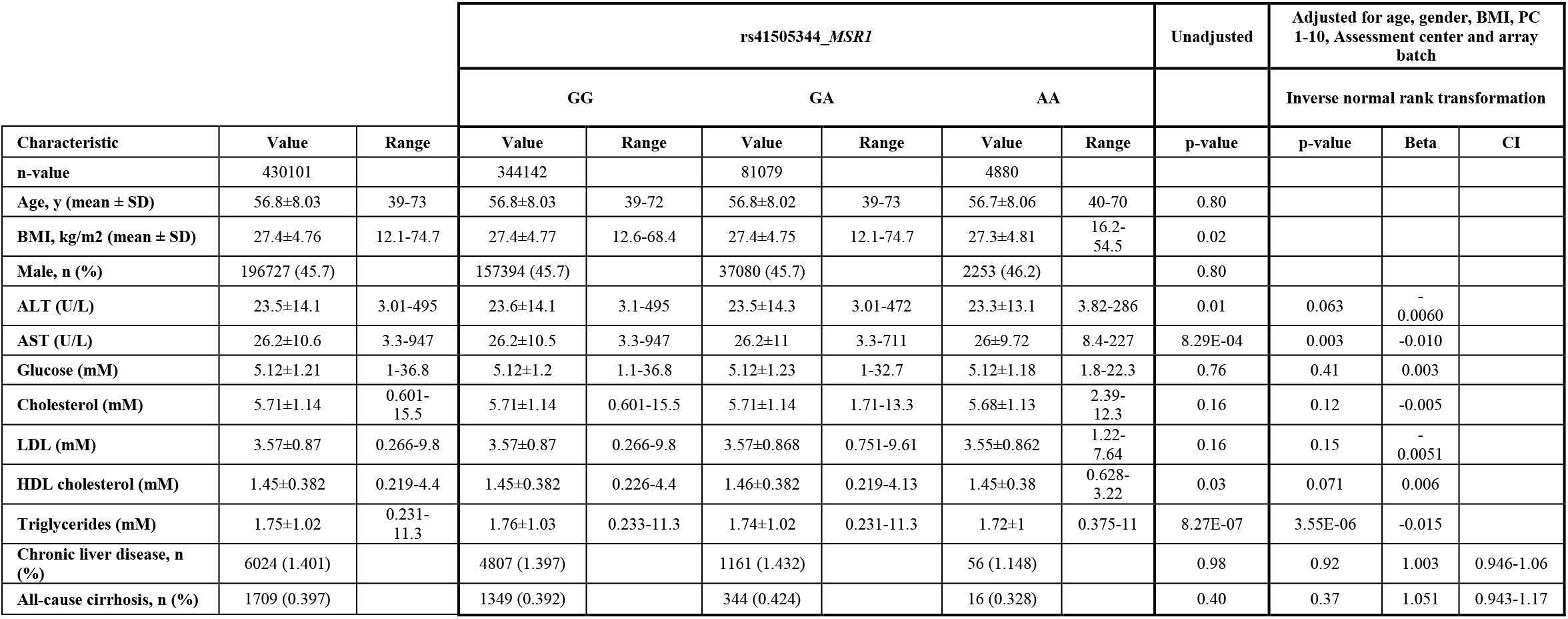
Correlation rs41505344 SNP with clinical features using UK Biobank (n=430,101)

Our human immunohistochemistry data indicated that MSR1 is expressed in the liver on mature endogenous macrophages, the Kupffer cells, rather than on infiltrating monocyte-derived macrophages. To unravel transcriptional regulatory mechanisms of *MSR1*, we used publicly available RNA sequencing data comparing human monocytes with differentiated macrophages, which identified 1,208 differentially expressed genes, with *MSR1* mRNA expression increased in the macrophage population (*24*). By motif enrichment analysis using iRegulon, we identified eight differentially expressed transcription factors, upregulated in human macrophages compared to monocytes, that are predicted to regulate the expression of *MSR1: BHLHE41, ETV5, HMGN3, MAF, MITF, NR1H3, THRA* and *ZNF562* (False discover rate<0.001, **Fig.6b**, **Table s4**). To verify whether these transcription factors bind any regulatory regions near the *MSR1* gene, and in particular the rs41505344 SNP locus, we investigated ChIP-sequencing data for these proteins. *MITF, MAF, THRA* and *NR1H3* proved to bind in the vicinity of the rs41505344 locus, suggesting an indirect role for the SNP in the transcriptional regulation of *MSR1* (**Fig.6c**). Taken together, these results suggest there is an increased frequency in NAFLD of variants potentially affecting *MSR1* expression during monocyte-macrophage differentiation, which thereby could influence features of obesity-related diseases.

## Discussion

In this study, we provide evidence that MSR1 is important for the uptake of lipids in macrophages leading to an inflammatory response and metabolic changes throughout the body. In a setting of lipid overload, MSR1 deficiency not only led to reduced hepatic inflammation and changes in hepatic lipid metabolism but it also reduced circulating fatty acids, increased lipid storage in the adipose tissue and improved glucose tolerance, highlighting the importance of the liver-adipose tissue axis in NAFLD and the metabolic syndrome (*25*). Our data demonstrated that in NAFLD biopsies MSR1 expression was observed in tissue-resident macrophages, the Kupffer cells, rather than in infiltrating monocytes, located in the portal tract (*26*). Previous research showed that differentiation of human monocytes towards mature macrophages goes hand-in-hand with increased expression of *MSR1* mRNA (*24*). The association between *MSR1* mRNA and features of steatohepatitis (hepatocyte ballooning and lobular inflammation) as found in our data, would suggest that there is an ongoing differentiation from infiltrating monocytes towards macrophages during NASH. Chronic portal inflammation is a histological feature correlated with advanced NAFLD in both adults and children (*27, 28*), yet it has not been associated with lobular inflammation, which is a predictor of fibrosis progression in patients with NAFLD (*29*). This would suggest that steatohepatitis is driven by mature tissue-resident macrophages rather than infiltrating monocytes. Our results support this as *Msr1* deficiency in HFD-fed mice tempered the lipid-induced inflammatory response in the liver, by reducing the expression of *Axl, Il1b, S100a8/a9* and *Spp1* but also *Cd44*. Cd44 expression has been associated with NASH in human and mouse, and is crucial for homing of monocytes into the damaged liver, suggesting that lipid accumulation in tissue-resident macrophages via MSR1 is a trigger to recruit immune cells (*30*). This is in line with previous reports where it was described that Kupffer cell depletion by clodronate liposomes in mice on a choline-deficient l-amino acid-defined diet for 22 weeks suppresses the infiltration of inflammatory cells, mainly monocytes, into the liver (*31*). Furthermore, our results showed that the absence of Msr1 induced a change in hepatic expression of genes associated with lipid metabolism, including an increase in *Ppara*, reduced hepatic triglycerides with concordantly increased mitochondrial oxygen consumption and ameliorated glucose tolerance in HFD-fed mice. Peroxisome proliferator-activated receptors (PPAR) are nuclear receptors playing key roles in metabolic homeostasis and inflammation (*32*). Selective depletion of Kupffer cells has been reported to activate Ppara signaling in hepatocytes while resulting in overall reduced levels of hepatic triglycerides in mice fed a 45%-HFD for 20 weeks (*33*). Furthermore, hepatocyte-restricted *Ppara* deletion in mice impaired liver lipid metabolism, leading to increased hepatic triglycerides and increased plasma FFAs (*34*). In human adult non-cirrhotic NASH patients, the PPAR agonist lanifibranor proved to induce NASH resolution and fibrosis regression after 24 weeks of treatment in a Phase 2b randomized, placebo-controlled, double-blind study (*35*). Taken together, the effect of Msr1 deficiency we have observed in this study on liver metabolism, triglycerides and circulating FFAs could in part be explained by the changed Ppara signaling in the liver.

In this study, we showed that MSR1 can facilitate the uptake of SFAs, such as palmitic acid, as well as non-SFAs, such as oleic acid, independent from other receptors. Yet, only SFAs could induce the release of TNFa through phosphorylation of JNK in macrophages, which is in line with previous reports (*12–15*). In murine methionine-and choline-deficient diet models, jnk1, but not jnk2, proved to be critical for diet-induced JNK activation and the development of steatohepatitis (*36*). In our *Msr1*^-/-^ HFD-fed mice, we observed lower *Tnfa* expression in the liver and adipose tissue as well as lower serum Tnfa levels. Furthermore, we showed that therapeutic blocking of MSR1 *in vivo* or *ex vivo* using human liver slices prevented the formation of foamy macrophages and reduced the release of TNFa. TNFa is of clinical relevance as serum levels have been reported to be increased in obese patients and correlated with NAFLD disease activity (*37, 38*). The effect of TNFa is pleiotropic as it can sensitize hepatocytes to apoptosis but it can also stimulate hepatic lipid synthesis and reduce *Ppara* expression (*39–41*). Furthermore, Tnfa affects glucose homeostasis in adipocytes and promotes lipolysis in cultured adipocytes (*42*).

Although current efforts to develop drug therapies for NAFLD primarily focus on ameliorating the specific histological features of the disease (i.e. steatohepatitis or fibrosis), it is important to remember that NAFLD is part of a multi-system metabolic disease state and so agents that offer more broad metabolic or cardiovascular benefits would be highly attractive (*43*). Our data indicate that by targeting MSR1, one would not only reduce lipid-induced inflammation in the liver but also improve dyslipidemia and affect improved lipid storage in the adipocytes. In addition, we demonstrate the feasibility of using targeted monoclonal antibody therapy to treat NASH by reducing hepatic inflammation *in vivo* in mice and *ex vivo* in human models. Moreover, we found some evidence that the genetic variant rs41505344 in *MSR1* was associated with serum triglycerides and ALT in a large cohort of over 400,000 patients. Though the SNP in *MSR1* was not strongly associated with susceptibility to NAFLD, we found that several transcription factors regulating the expression of *MSR1* bound in the locus of this SNP. This would suggest that rs41505344 could influence the expression of *MSR1* during macrophage differentiation/maturation and hence effect levels of serum triglycerides and ALT.

There are several limitations to this study. Although we have used large cohorts of adult patients, we do not know if we do not know if the results would be similar in paediatric NAFLD. In addition, we did not provide any functional proof for the role of rs41505344 in a setting of lipid-overload. To investigate the effect rs41505344 on *MSR1* expression and features of the metabolic syndrome, a knock-in mouse model challenged with a long-term diet could help to provide answers. Moreover, in this current study we used a global knock-out mouse model and focused on the early phases of NAFLD by using a relative short-term diet of 16 weeks. To further investigate the liver-adipose tissue axis, a Kupffer cell-specific *Msr1* knock-out or a conditional *Msr1* knock-out mouse model could be used. In this study, we focused on MSR1 in a setting of lipid overload, yet other macrophage scavenger receptors have been reported to play a role during the metabolic syndrome. TREM2 has been described to induce liver fibrosis and regulate adipose tissue homeostasis (*19, 20*), while soluble CD163 has been associated with adipose tissue insulin resistance, circulating FFAs and hepatic steatosis (*44*). Although, our RNA sequencing data of HFD-fed *Msr1*-/- mice showed no differential expression for these receptors, this does not necessarily exclude synergy in function or pathway signaling between these receptors.

This study showed that the scavenger receptor MSR1, as part of the innate immune system, is a critical sensor for lipid homeostasis, highlighting the importance of the liver-adipose tissue axis. With the prevalence of obesity increasing globally, it is crucial that we understand how our immune system reacts when challenged with over-nutrition. Understanding and therapeutically influencing macrophage immunometabolism, could help us treat features of the metabolic syndrome, such as dyslipidemia, NAFLD and type II diabetes.

## Materials and Methods

### Study design

A total of 170 adult NAFLD liver biopsies were processed for transcriptomic analysis and MSR1 protein expression was assessed in 14 liver and 31 adipose tissue biopsies, and correlated with clinicopathological features. To functionally validate our findings, *Msr1*-/- and WT mice were submitted to a high fat and cholesterol diet for 16 weeks. The Msr1 receptor and down-stream signaling targets were inhibited *in vitro* using isolated mouse primary macrophages and bone marrow-derived macrophages, challenged with lipids. Therapeutic intervention with monoclonal antibody against MSR1 was performed in WT mice on a 12 week high fat and cholesterol diet, and in lipid loaded *ex vivo* human liver slices. To assess susceptibility with NAFLD and the metabolic syndrome, GWAS analysis was performed using previously published data including 1,483 European NAFLD cases and 17,781 European general-population controls (*23*). Results were replicated using clinical data from 430,101 participants of the UKBiobank. Publicly available RNA sequencing data was used to investigate the association between SNPs and transcription factors regulating *MSR1* expression when differentiating human monocytes towards macrophages.

### Patient selection

Cases were derived from the European NAFLD Registry (*45*). For the histopathological and nanoString^®^ study, 215 liver and adipose tissue samples from Caucasian patients were included. 184 formalin-fixed paraffin-embedded (FFPE) or frozen liver biopsies samples were obtained from patients diagnosed with histological proven NAFLD at the Freeman Hospital, Newcastle Hospitals NHS Foundation Trust, Newcastle-upon-Tyne, UK and at the Pitié-Salpêtrière Hospital, Paris, France. In addition, 20 FFPE subcutaneous adipose tissue samples and 11 matching omental adipose tissue samples were obtained from obese patients with or without NAFLD, diagnosed and treated at the Pitié-Salpêtrière Hospital, Paris, France (Table S5). For the Genome Wide Association Study, 1,483 patients with histological proven NAFLD recruited at different European centers were included as previously described (*23*). All liver tissue samples for the histopathological and nanoString^®^ study were centrally scored according to the semiquantitative NASH-CRN Scoring System by an expert liver pathologist (DT) (*18*). Fibrosis was staged from F0 through to F4 (cirrhosis). Alternate diagnoses and etiologies such as excessive alcohol intake, viral hepatitis, autoimmune liver diseases and steatogenic medication use were excluded. Viable human normal liver tissue for the *ex vivo* slices was obtained after resection from two adult patients treated at the University Hospitals Leuven, Leuven, Belgium. Samples were assessed by an expert liver pathologist (TR). This study was approved by the local or national relevant Ethical Committees of each in the participating center.

### nanoString

mRNA was isolated from 170 liver biopsy samples (FFPE and frozen). AllPrep DNA/RNA Micro kit (Qiagen) was used for the frozen samples, the High Pure FFPET RNA Isolation Kit (06650775001, Life Science Roche) for the FFPE biopsies. Custom-made assay panels were run on the nanoString^®^ nCounter analysis system (nanoString^®^). The nSolver 3.0 software (nanoString^®^) was used for positive and negative control probe validation and the data were normalized to three housekeeping genes (*RPL19, SRSF4* and *YWHAZ*).

### Animals

*Msr1*-/- C57BL/6 mice were kindly provided by Prof. Siamon Gordon, University of Oxford. All mice were maintained under specific pathogen free conditions and experiments were approved by the Institutional Animal Care and Use Committee and with a project license of the UK Home office. Mice had free access to water and were fed either standard chow (n=10, 5 WT and 5 *Msr1*-/-) or high-fat and high-cholesterol diets (n=10, 5 WT and 5 *Msr1*-/-, 45%-HFD; 820263, Special Diet Services) ad libitum. Body weight was measured weekly during the feeding period and compared with liver weight and visceral fat mass after the animals were sacrificed. For the therapeutic intervention study, WT mice were put on a high-fat and high-cholesterol diet (45%-HFD; 820263, Special Diet Services) for 12 weeks and treated with monoclonal rat anti-mouse Msr1 antibody (n=3 animals, (MAB1797-SP) R&Dsystems) or rat IgG control (n=4 animals, (MAB0061) R&D systems) at week 10 and 11 by intravenous injection (0.25 mg of Msr1 antibody or corresponding IgG control for each animal).

### Statistical analysis

Data were tested for normality using the Kolmogorov-Smirnov or the Shapiro-Wilk normality test using GraphPad Prism 8.0.1 (GraphPad Software Inc.). Unpaired Student’s t-test or Mann-Whitney U test, one way ANOVA or Kruskal-Wallis test with respectively Tuckey’s or Dunn’s post hoc multiple comparison test, and Spearman’s rho correlation were performed using GraphPad Prism 8.0.1. Data were visualized using GraphPad Prism 8.0.1. A p-value<0.05 was considered significant.

*Additional Material and Methods can be found in the Supplementary Materials*.

## Supplementary Materials

Fig. S1. Macrophage Scavenger Receptor 1 (MSR1) expression in human non-alcoholic fatty liver disease (NAFLD)

Fig. S2. Expression of MSR1 and CD68 in human NAFLD

Fig. S3. Characterization HFD-fed *Msr1*-/- mice.

Fig. S4. Co-cultures of WT and *Msr1*^-/-^ BMDMs with adipocyte-and hepatocyte-like cells.

Fig. S5. Msr1 regulates lipid-induced pro-inflammatory activation of macrophages through JNK signaling.

Table S1. Demographics patient cohort used for nanoString analysis.

Table S2. Differentially expressed genes comparing HFD-fed *Msr1*-/-with WT mice using RNA sequencing.

Table S3. Top *MSR1* SNPs in GWAS analysis NAFLD Case (n=1,483) vs control (n=17,781) with p<1.0E-04

Table S4. iRegulon analysis identifying upregulated transcription factors binding to MSR1 region when comparing macrophages with monocytes.

Table S5. Clinical data obese patient cohort used for immunohistochemistry.

## Acknowledgments

The authors would like to thank the Newcastle Bioimaging Unit, the Newcastle University Genomics Core Facility, the Newcastle NanoString Core Facility and the Newcastle Molecular Pathology Node Proximity Laboratory for their technical support.

## Author contributions

OG and AH conceived the study. Study design, manuscript drafting and funding: AH, OG, MT and QMA. Manuscript preparation: AH, OG, SKP, MT, QMA. *In vivo* experiments: AH, SKP, OBG and NML. *In vitro* experiments: AH, OG and SKP. Human *ex vivo* experiments: OG, MVH, TR. Histopathology: OG, MVH, TR and DT. Nanostring analysis: OG. Bioinformatics: OG and JW. GWAS analysis: RD, HJC, AKD. eQTL UKBiobank data: RMM, OJ, SR. All authors contributed to data collection and interpretation, and critically revised the manuscript for intellectual content.

## Data and materials availability

To review GEO accession GSE163471:

Go to https://www.ncbi.nlm.nih.gov/geo/query/acc.cgi?acc=GSE163471

